# Network Biomarkers of Alzheimer’s Disease Risk Derived from Joint Volume and Texture Covariance Patterns in Mouse Models

**DOI:** 10.1101/2025.02.05.636582

**Authors:** Eric W. Bridgeford, Jaewon Chung, Robert J. Anderson, Ali Mahzarnia, Jacques A. Stout, Hae Sol Moon, Zay Yar Han, Joshua T. Vogelstein, Alexandra Badea

## Abstract

Alzheimer’s disease (AD) lacks effective cures and is typically detected after substantial pathological changes have occurred, making intervention challenging. Early detection and understanding of risk factors and their downstream effects are therefore crucial. Animal models provide valuable tools to study these prodromal stages. We investigated various levels of genetic risk for AD using mice expressing the three major human APOE alleles in place of mouse APOE. We leverage these mouse models utilizing high-resolution magnetic resonance diffusion imaging, due to its ability to provide multiple parameters that can be analysed jointly. We examine how APOE genotype interacts with age, sex, diet, and immunity to yield jointly discernable changes in regional brain volume and fractional anisotropy, a sensitive metric for brain water diffusion. Our results demonstrate that genotype strongly influences the caudate putamen, pons, cingulate cortex, and cerebellum, while sex affects the amygdala and piriform cortex bilaterally. Immune status impacts numerous regions, including the parietal association cortices, thalamus, auditory cortex, V1, and bilateral dentate cerebellar nuclei. Risk factor interactions particularly affect the amygdala, thalamus, and pons. APOE2 mice on a regular diet exhibited the fewest temporal changes, suggesting resilience, while APOE3 mice showed minimal effects from a high-fat diet (HFD). HFD amplified aging effects across multiple brain regions. The interaction of AD risk factors, including diet, revealed significant changes in the periaqueductal gray, pons, amygdala, inferior colliculus, M1, and ventral orbital cortex. Future studies should investigate the mechanisms underlying these coordinated changes in volume and texture, potentially by examining network similarities in gene expression and metabolism, and their relationship to structural pathways involved in neurodegenerative disease progression.

## 1 Introduction

Alzheimers disease (AD) is the most common type of dementia and is estimated to affect more than 5 million U.S. citizens and more than 25 million people worldwide. The risk of Alzheimer’s disease (AD) is complex and multifactorial, resulting in multiple pathologies, and is influenced by factors including genetic predispositions [1], environmental factors [2], lifestyle [3], and most importantly age [4]. Therefore, identifying and understanding which regions of the brain are highly vulnerable, that is, subject to significant changes, and how different risk factors contribute to this vulnerability, is important to understand, bearing the potential to open new therapeutic targets and opens new horizons for AD prevention [5].

Our scientific premise is based on the role of alleles of the apolipoprotein E (ApoE) gene in aging and Alzheimer’s disease. ApoE is a critical gene involved in lipid metabolism and neuronal repair processes, with its variants: ApoE2, ApoE3, and ApoE4 playing distinct roles in neurodegenerative diseases [6, 7]. The most studied among these, ApoE4, is known for its strong association with an increased risk of Alzheimer’s disease. In contrast, ApoE2 is considered neuroprotective, while ApoE3 is neutral. This genetic variability offers a unique opportunity to explore the connection between genetic profiles and brain structure, particularly in relation to neurodegenerative diseases. Mouse models with human targeted replacement APOE alleles allow for understanding the mechanisms behind early alterations in such association at prodromal stages.

Recent research in AD is increasingly focused on understanding the relationship between brain volume variations, also known as structural covariance/correlation. A notable study demonstrated that single nucleotide polymorphisms (SNPs) susceptible to Alzheimer’s disease are closely related to changes in grey matter volume and cognitive outcomes [8]. This research utilized magnetic resonance imaging to construct grey matter structural covariance networks (SCNs) in patients with Alzheimer’s disease. The study assessed the effects of various genetic loci on cognitive outcomes, revealing that specific genetic variations, including the ApoE4 allele, interact with or independently affect cognitive outcomes. Other studies similarly support the fact that there is a relationship between AD progression and changes in SCNs [8, 9]. However, these methods exhibit limitations. The main limitation is that these methods construct a single SCN per group, which can lead to a small sample size when trying to find brain regions that are changing, rather than a single correlation between pairs of regions. Performing hypothesis tests on individual elements of a correlation matrix, whose data are highly correlated, can result in spurious conclusions if the interactions between regions are not decoupled.

To address these issues, we propose a method for obtaining individualized representations of covariance and leverage recent advances in *K*-sample and multi-way hypothesis testing framework to identify brain regions that are highly vulnerable, or susceptible to significant change [10]. We first compute an absolute difference of a feature between all pairs of brain regions, resulting in a network per subject. We focus on studying the brain volume derived from structural MRI, and fractional anisotropy derived from diffusion MRI. Given these networks, we obtain a low-rank representation for each brain region per subject by jointly modeling the networks. We then leverage distance correlation to perform multi-way tests that include factors such as age, sex, genotypes, and diet.

We used magnetic resonance microscopy [11] to derive quantitative metrics reflective of brain microstructural integrity in mouse models with different risk actors for AD (age, ApoE genotype, sex, diet and immunity). Specifically, we employed diffusion imaging, since this can reveal multiple markers for both volume and microstructural changes [12]. Our results reveal significant influence of genetic risk factors, particularly in differentiating between amygdala network-associated and non-associated ApoE allelic populations. While alone age, sex, and diet emerge as marginal factors, their significance escalates when considering the broader ApoE allelic set. These insights not only improve our understanding of ADs complexity but also signify a pivotal step towards stratified, personalized and effective clinical interventions.

**Figure 1:**
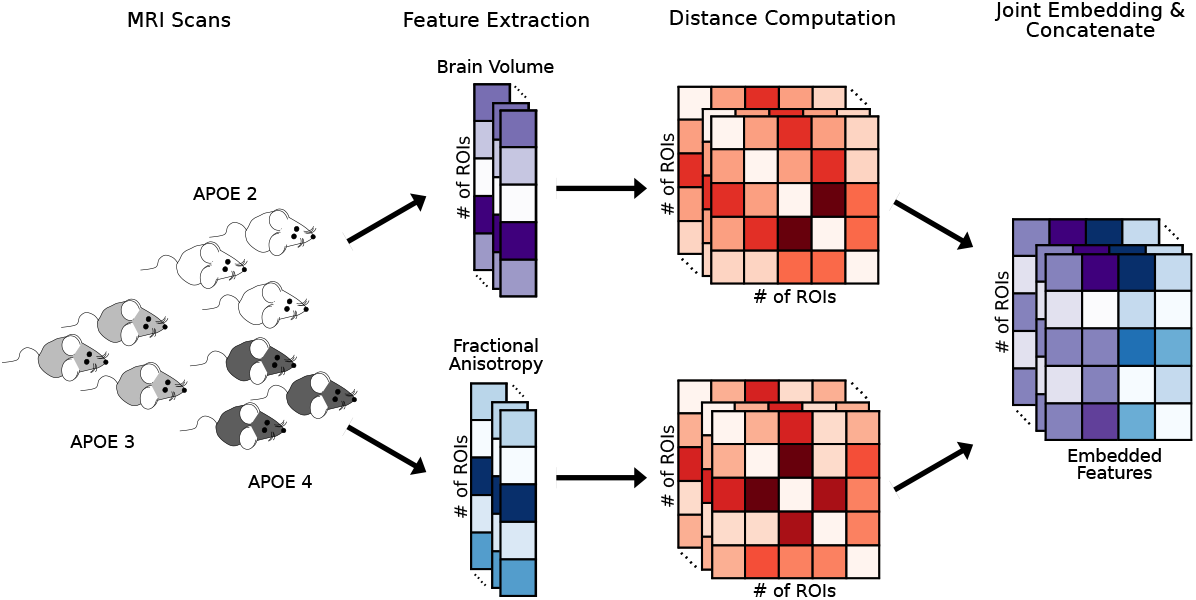
An illustration of the preprocessing pipeline. MRI scans from the three genotypes of mice, ApoE2, ApoE3, and ApoE4 are processed to estimate brain volume and fractional anisotopy data per ROI. Each data modality is then transformed into distance matrices, which are jointly embedded. Resulting embedded features are concatenated for subsequent analysis.

## 2. Materials and Methods

### 2.1 Animals

We have used mice expressing the human apolipoprotein E (ApoE) alelles, critically involved in lipid metabolism and neuronal repair processes [13], homozygous for its variants–ApoE2, ApoE3, or ApoE4–because of their distinct roles in neurodegenerative diseases. APOE3 is the allele found in a majority of the population. The most studied among these, ApoE4, is known for its strong association with an increased genetic risk of Alzheimer’s disease. In contrast, ApoE2 is considered neuroprotective [14]. All mice have the human version of the ApoE allele, and some express the mouse nitric oxide synthase gene (Nos2) while others express the human nitric oxyde synthase 2 (NOS2) gene (HuNOS2tg/mNos2^*−/−*^) mice, termed HN).

This mutation helps address differences between the human and mouse inflammatory responses, where human macrophages, express little NOS2, and generate much less nitric oxide (NO) in response to inflammatory stimuli, compared to mouse macrophages [15]. Introducing the human NOS2 gene lowers the amounts of NO produced, to help render the innate immune system more human like by bringing the mouse immune/redox activity more in tune with the human [16].

Mice were fed a control diet for their whole life (2001 Lab Diet; denoted Ctrl), or swithced to a high fat diet for the last 4 months before sacrifice (D12451i, Research Diets; denoted HFD). A total of 169 mice were scanned using diffusion weighted MRI (dMRI). See Table 1 for more details.

**Table 1:** Description of the mice population. For each genotype, some mice were given high fat diet (HFD) and some had the humanized NOS2 gene.

| Genotype | N | Sex (Female) | Age (Mean; Std) | High Fat Diet | Immunity |
| --- | --- | --- | --- | --- | --- |
| ApoE 2 | 58 | 30 | 16.12; 1.8 | 15 | 25 |
| ApoE 3 | 54 | 27 | 15.25; 1.54 | 25 | 20 |
| ApoE 4 | 57 | 28 | 15.84; 1.76 | 15 | 30 |

Specimens were prepared for imaging as described in [17]. Briefly mice were anesthetized with ketamine/xylazine (100 mg/kg ketamine, 10 mg/kg xylazine) and perfused via the left cardiac ventricle. Blood was cleared with 0.9% Saline at 8 ml/min for 5 min. Fixation was done perfusing 10% neutral buffered formalin phosphate containing 10% (50 mM) Gadoteridol (ProHance, Bracco Diagnostics Inc., United States) at 8 ml/min for 5 min. The specimens were then stored in formalin for 12 h, then moved in phosphate-buffered saline with 0.05% (2.5 mM) Gadoteridol until imaging. Specimens were imaged in the skull, to avoid brain damage and distortion, and while submerged in fomblin to reduce susceptibility artifacts and prevent dehydration.

### 2.2 Image Collection and Preprocessing

Mouse brain specimens were imaged at 9.4 T, as described in [18], using a 3D spin echo diffusion weighted imaging (SE DWI) sequence with TR/TE: 100 ms/14.2 ms; matrix: 420 *×* 256*×* 256; FOV: 18.9 *×*11.5 *×* 11.5 mm, BW 62.5 kHz; reconstructed at 45 µm isotropic resolution. Diffusion weighting was applied along 46 directions, using 2 diffusion shells (23 directions using a b value of 2,000 s/*mm*^2^ and 23 directions using a b value of 4,000 s/*mm*^2^); we also acquired 5 non-diffusion weighted images (b0). The max diffusion pulse amplitude was 130.57 Gauss/cm; duration 4 ms; separation 6 ms. We used eightfold compressed-sensing acceleration and reconstructed images using BART [19] [20].

Diffusion parameters were reconstructed using MRtrix3 [21] producing 2 million tracts per brain. To produce regional estimates of volume and FA we have used pipelines implemented in a highperformance computing environment for image reconstruction [22], and atlas based segmentation [23, 24] using a symmetrized mouse brain atlas [25, 26] with 332 regions, 166 for each hemisphere.

### 2.3 Single Subject Networks and Covariates

Each mouse brain image was segmented into 332 brain regions using SAMBA, and the symmetrized mouse brain atlas previously used for connectomic analyses [23, 24]. The volume of each brain region was computed by counting the number of voxels in each region and multiplying this by the voxel size, and the average fractional anisotropy (FA) for each brain region was calculated using MRtrix3[21] (see Section 2.1 and 2.2 for more details).

To compute single-subject networks, we first normalized brain volume and FA by dividing by the maximum brain volume and FA, respectively, then computed the absolute difference between all pairs of brain regions, resulting in a distance matrix of size 332 *×* 332, where each element of the matrix corresponds to the difference of either brain volume or FA from a pair of brain regions. This resulted in two networks per mouse, one from each of the two data modalities. For all subsequent analyses, we treated these networks as undirected (since we computed absolute differences), weighted so that all values lie between 0 and 1, and loopless (e.g. there is no difference in volume of a particular region to itself). Figure 2A and 2C shows the averaged distance matrices for brain volume and FA, respectively. To better understand the distance matrices, we then computed the difference of the distance matrices from pairs of genotypes, which highlights qualitative differences in the distance matrices (Figure 2B and 2D).

**Figure 2:**
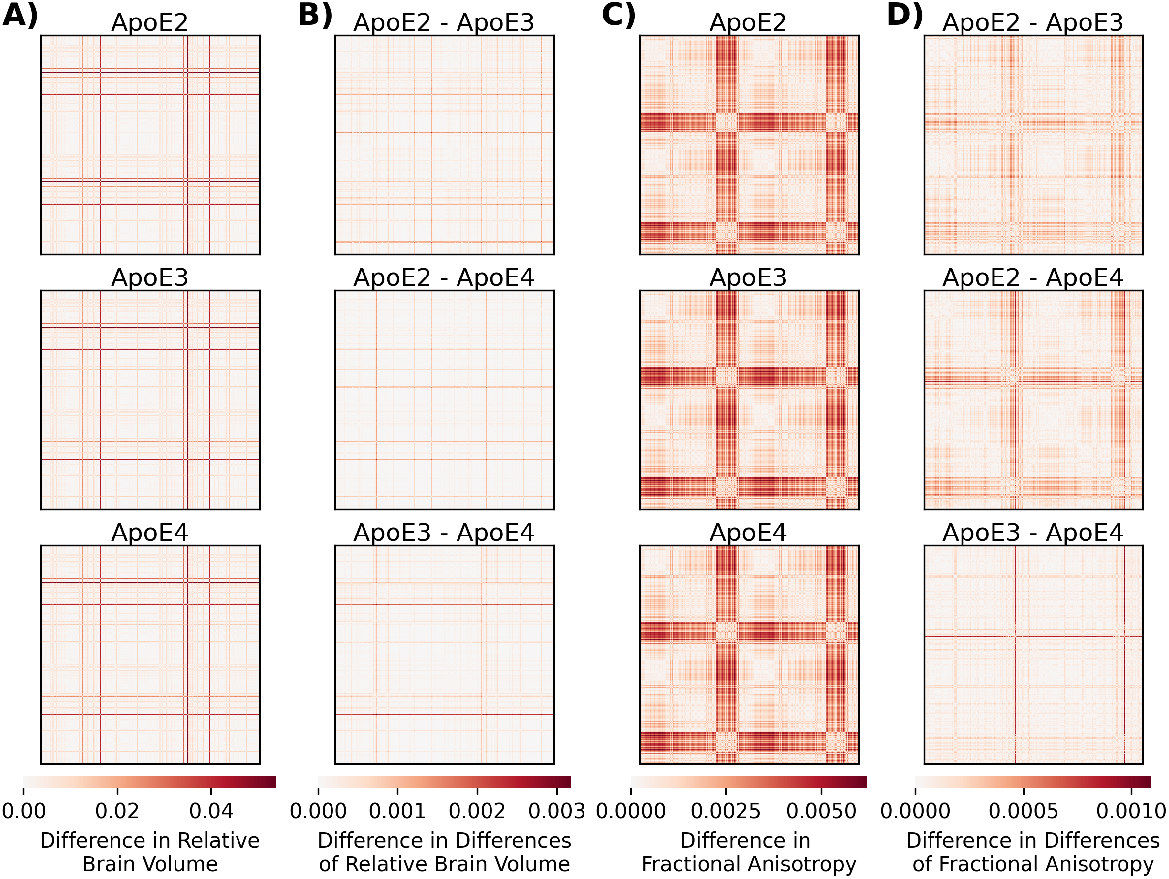
Visualization of the Distance Matrices from Brain Volume (BV) and Fractional Anisotropy (FA) Measurements. **(A)** The averaged difference of **brain volumes** between all pairs of brain regions. Each row represents one of the three ApoE genotypes. **(B)** The difference in differences of relative brain volume between each possible pairs of three ApoE genotypes. **(C)** The averaged difference of **fractional anisotropy** between all pairs of brain regions. Each row represents one of the three APOE genotypes. **(D)** The difference in differences of fractional anisotropy between each possible pairs of three ApoE genotypes.

Besides brain volume and fractional anisotropy (FA) data, we collected additional information for each mouse that might predict Alzheimer’s disease risk. This included: genotype (ApoE2, ApoE3, or ApoE4, which have varying risk levels), sex (female or male, as females are believed to be more susceptible), age (categorized as below or above the median age for each genotype, since older age is a major risk factor for late-onset AD), immunity (presence of the humanized NOS2 gene, denoted as HN, for a more human-like immune system), and diet (control or high-fat diet (HFD), with HFD expected to increase vulnerability).

Next, we present a model for multiple networks and methods for obtaining new representations of all the networks, which takes into account the inherent structure and dependencies within networks.

### 2.4 Statistical Modeling of Networks

Statistical models for networks allows us to model all of the inherent dependencies across vertices (brain ROIs) in networks, and transforms the data in which we can apply traditional statistical and machine learning tools that would otherwise be inappropriate for network data. The joint random dot product graph (`JRDPG`) model provides a way to model weighted networks and allow us to obtain a Euclidean representation of networks using statistically principled procedures [27–29]. In this model, a vertex is a region-of-interest (ROI) in the brain, which is represented as a low-dimensional vector called a latent position. The probability of one ROI connecting to another is determined by the dot product of the corresponding latent positions. In other words, a matrix containing the latent positions of all ROIs is a representation of the underlying distribution of the networks.

Given our networks derived from brain volume and FA data, we can obtain the low-dimensional latent positions using a joint embedding technique called omnibus embedding (`Omni`) (see Section 2.6 for more details); that is embed all of the networks from derived from brain volume and then embed networks from the FA independently. We will then concatenate the output embeddings for each modality giving us a final feature set that will be used for subsequent analysis. The dimensionality of the latent position is determined by an automatic elbow detection algorithm [30], which is 3 for both brain volume and FA. We finally obtain a 6 dimensional latent position vector per brain region, resulting in a matrix of size 332 *×* 6 per mouse, which we will use for our subsequent analysis. Next, we detail a framework for understanding and quantifying the differences in these maps and how we construct a hypothesis test using latent positions and predictor variables.

### 2.5 Hypothesis Testing for Discovering Vulnerable Regions

We sought to understand whether these single-subject maps were significantly different according to some definition, in order to identify and characterize how certain predictor variables, such as ApoE genotypes, affect brain regions. A simple approach is to ask whether the brain volume and FA of a region are significantly different across a predictor. One way to formalize the question into a hypothesis test is as a *K* -sample testing problem. Let *y*^(*i*)^ represent measurements (possibly vector-valued high-dimensional) for *i* = 1, …, *m* samples, where for each sample, we have additional covariates *t*^(*i*)^ where are from one-of-*K* groups.

For example, *t*^(*i*)^ can denote different genotypes such as ApoE2, ApoE3, and ApoE4, and *y*^(*i*)^ can be measurements, such as brain volume and FA. Assume that if *t*^(*i*)^ = *k*, that measurements *y*^(*i*)^ are sampled independently and identically from some distribution *F*_*k*_. A reasonable test would be whether the measurements differ depending on the grouping; this can be formalized as the *K* -sample test:

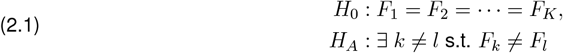

In words, under the null hypothesis, the measurements do not differ across groups; under the alternative, there exists at least one pair of groups for which the measurement distributions are unequal. Intuitively, rejection of the null hypothesis in favor of the alternative has the interpretation that the grouping encodes meaningful differences in the measurements for a particular region (the region is vulnerable). This test can be generalized to the case where there are multiple predictors, or multi-way *K*-sample testing. For example, suppose that *K* denotes our three ApoE genotypes and that for each sample we obtain additional predictor data, such as age and sex. Traditionally, analysis of variance (`Anova`) [31] or multivariate `Anova` (`Manova`) [32] are conventional choices for *k* -sample tests. For multi-way *K*-sample testing, *K* -way `Anova` [33] or *K* -way `Manova` are common choices. However, these tests often do not perform well for high-dimensional or non-Gaussian data, which is typically the case for network data, because their performance depends on assumptions that are generally not present in real data [34, 35]. Several non-parametric alternatives have been developed to address this issue; we choose distance correlation (`DCorr`) for testing [36]. While `DCorr` is an independence test, the connections between the independence test and the multi-way *K*-sample test are detailed in Panda et al. [10].

To begin our investigation, we divided the analysis into two distinct categories due to the absence of observed data for certain combinations of factors. For instance, not all mice with a non-HN immunity gene were subjected to a high-fat diet. Consequently, we conducted two separate analyses to understand the impact of human immunity and diet. The first analysis examined the effects of genotypes, sex, age, and immunity (HN and non-HN), while the second analysis explored the effects of genotypes, sex, age, and diet (control and HFD). Specifically, only mice that were given normal diet were considered in the first analysis, and only mice with HN immunity were considered in the second analysis.

### 2.6 Graph Theory Preliminaries

Networks (or graphs) are convenient mathematical objects for representing connectomes. A network *G* consists of the ordered pair (*V, E*), where *V* is the set of vertices and *E* is the set of edges. The set of vertices can be represented as *V* = *{*1, 2, …, *n }*, where |*V* |= *n* is the number of vertices. The set of edges is a subset of all possible connections between vertices (i.e., *E ⊆ V ×V*). We say tuple (*i, j*) *∈ E* if there exists an connection between vertex *i* and vertex *j*. In many connectomics datasets, edges have associated edge weights: these are real-valued numbers that encode quantitative information about a connection between two vertices.

### 2.7 Statistical Models

Statistical modeling of connectomics data enables the principled analysis of these high-dimensional, graph-valued data. Random graph models treat individual connectomes as random variables, enabling mathematical characterization of network structure and accounting for noise within and across observed samples. Treating connectomes as random network-valued variables sampled from these random graph models enables the formulation of hypothesis tests that can be used to identify connective differences at multiple levels across numerous phenotypic profiles.

## Random Dot Product Graph Model

The Random Dot Product Graph (`RDPG`) is a type of independent edge model. In this model, each element of an adjacency matrix is sampled independently from a Bernoulli distribution:

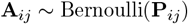

Given the number of vertices *n*, the matrix **P** is a *n × n* matrix of edge-wise connection probabilities with elements in [0, 1]. We can construct various models depending on the constraints imposed on **P**. Note that we assume that **P** has no self-loops (i.e. diag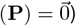and is undirected (i.e. **P**^*T*^ = **P**).

In the random dot product graph (`RDPG`), the probability of a connection **P**_*ij*_ is determined by the vertices. Each vertex *i ∈ V* is associated with a low-dimensional *latent position* vector, **X**_*i*_, in the Euclidean space ℝ^*d*^. The probability of connection between vertices *i* and *j* is given by the dot product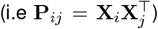. Thus, in a *d*-dimensional `RDPG` with *n* vertices, the rows of the matrix **X** ℝ^*n×n*^ are the latent positions of each vertex, and the matrix of edge-wise connection probabilities is given by **P** = **XX**^*T*^. Each element of the adjacency matrix is then independently modelled as

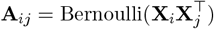

where **X**_*i*_ and **X**_*j*_ are latent positions for vertices *i* and *j*, respectively.

We acknowledge that the original intention of `RDPG` is to model binary networks, although the model can be naturally extended to handle weighted networks. However, the weighted `RDPG` models are not well studied, and as such does not enjoy the same statistical guarantees. In the subsequent section, we present an algorithm for estimating latent positions from observed data and describe methods for preprocessing the data to enable interpretation of results within the context of binary networks while still utilizing weighted network data.

### Joint Random Dot Product Graphs (JRDPG)

In this model, we consider a collection of *m* RDPGs all with the same generating latent positions. Similar to a RDPG, given an appropriately constrained Euclidean subspace ℝ^*d*^, the model is parameterized by a latent positions matrix **X** *∈* ℝ^*n×d*^ where *d ≪n*. The model is **A**^(1)^, **A**^(2)^, …, **A**^(*m*)^ *∼* JRDPG(**X**) where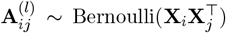 for all *i, j∈* [*n*] and *l∈* [*m*]. Each graph has marginal distribution **A**^(*l*)^ *∼*RDPG(**X**) for all *l∈* [*m*], meaning that the matrices **A**^(1)^, …, **A**^(*m*)^ are conditionally independent given **X** [37, 38].

### 2.8 Adjacency Spectral Embedding

The modeling assumptions of `RDPG` make the estimation of latent positions, which are usually unobserved in practice, analytically tractable. The estimation procedure we use is *adjacency spectral embedding* (`ASE`) [39]. The `ASE` of an adjacency matrix **A** in *d* dimensions is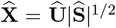, where|Ŝ|is a diagonal *d × d* matrix containing on the main diagonal the absolute value of the top-*d* eigenvalues of **A** in magnitude, in decreasing order, and Û is an *n × d* matrix containing the corresponding orthonormal eigenvectors. This simple and computationally efficient approach results in consistent estimates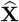 of the true latent positions **X** [39–41]. The `ASE` depends on a parameter *d* that corresponds to the rank of the expected adjacency matrix conditional on the latent positions; in practice, we estimate this dimension,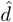, via the scree plot of the eigenvalues of the adjacency matrix which can be done automatically via a likelihood profile approach [30].

### Omnibus Embedding

Consider a sample of *m* observed graphs *𝒢* ^(1)^, *𝒢* ^(2)^, …, *𝒢* ^(*m*)^ and their associated adjacency matrices, **A**^(1)^, **A**^(2)^, …, **A**^(*m*)^ *∈* ℝ^*n×n*^ with *n* vertices that are identical and shared across all graphs. Under the JRDPG model, Omni is a consistent method for simultaneously estimating the latent position matrices for each graph by computing the spectral embedding into *d*-dimensions on the omnibus matrix, **O** *∈* ℝ^*nm×nm*^, as defined below

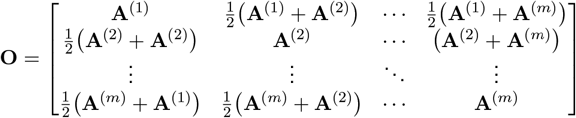

The embeddings gives the matrix

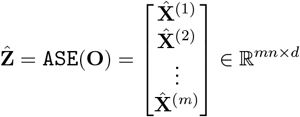

where the first *n* rows are the latent positions corresponding to **A**^(1)^, **A**^(2)^, …, **A**^(*m*)^.

#### Methodological intuition

Our procedures can be described intuitively as follows: under the null hypothesis (there is no difference in measurements across groups; e.g., Equation (2.1)), the JRDPG model is flexible for homogeneous networks, the Omni embedding provides an effective strategy for estimating the latent positions of these networks, and these estimated latent positions will have approximately the same distribution across groups. Under the alternative hypothesis, the latent positions estimated by Omni will differ; e.g., the estimated latent positions will not have the same distribution across groups. We exploit these observations to motivate the use of the estimated latent positions from volume and FA-derived difference matrices for our testing procedures described herein.

For our statistical analyses, we use the estimated latent positions for each ROI jointly across both volume and average fractional antisotropy to test if they differ significantly given some combination of groups (e.g. certain labels). For example, we test of there is a difference among the ApoE 2, 3, 4 genotypes while considering both sex (male and female), as well as immunity (presence or absence of HN). We then run subsequent analysis comparing subgroups. For example, instead of examining all three genotypes, we can compare APOE2 and 3 while considering gender and alleles.

### 2.9 p-values and Multiple Hypothesis Correction

All *p*-values from `DCorr` tests are estimated using *N* = 25,000 permutations via a chi-squared approximation for fast computations [42]. Across all figures and tables associated with this work, we are interested in interpreting individual statistical tests at a given significance level. Therefore, we control the familywise error rate (FWER) with the Holm-Bonferroni correction [43]

## 3 Results

### 3.1 Visualizations of Distance Matrices Show Significantly Different Regions

Distance matrices for volume and fractional anisotropy (FA) were qualitatively different, yet both supported differences between genotypes, with some consistency between pairwise genotype differences, but showed more widespread effects in FA texture

### 3.2 Effects of risk factors in animals on a control diet

Our multi-way nonlinear hypothesis testing addressed the influence of APOE genotype, immunity, sex, and age on the brain volume and FA variations within specific brain regions

In these comparisons, we only study mice with normal diet, and no mice with high fat diet were included as some mice with the non-humanized ApoE gene were treated with high fat diet. Figure 3 shows the statistical significance of each factor in various areas of the brain, which are summarized below.

**Figure 3:**
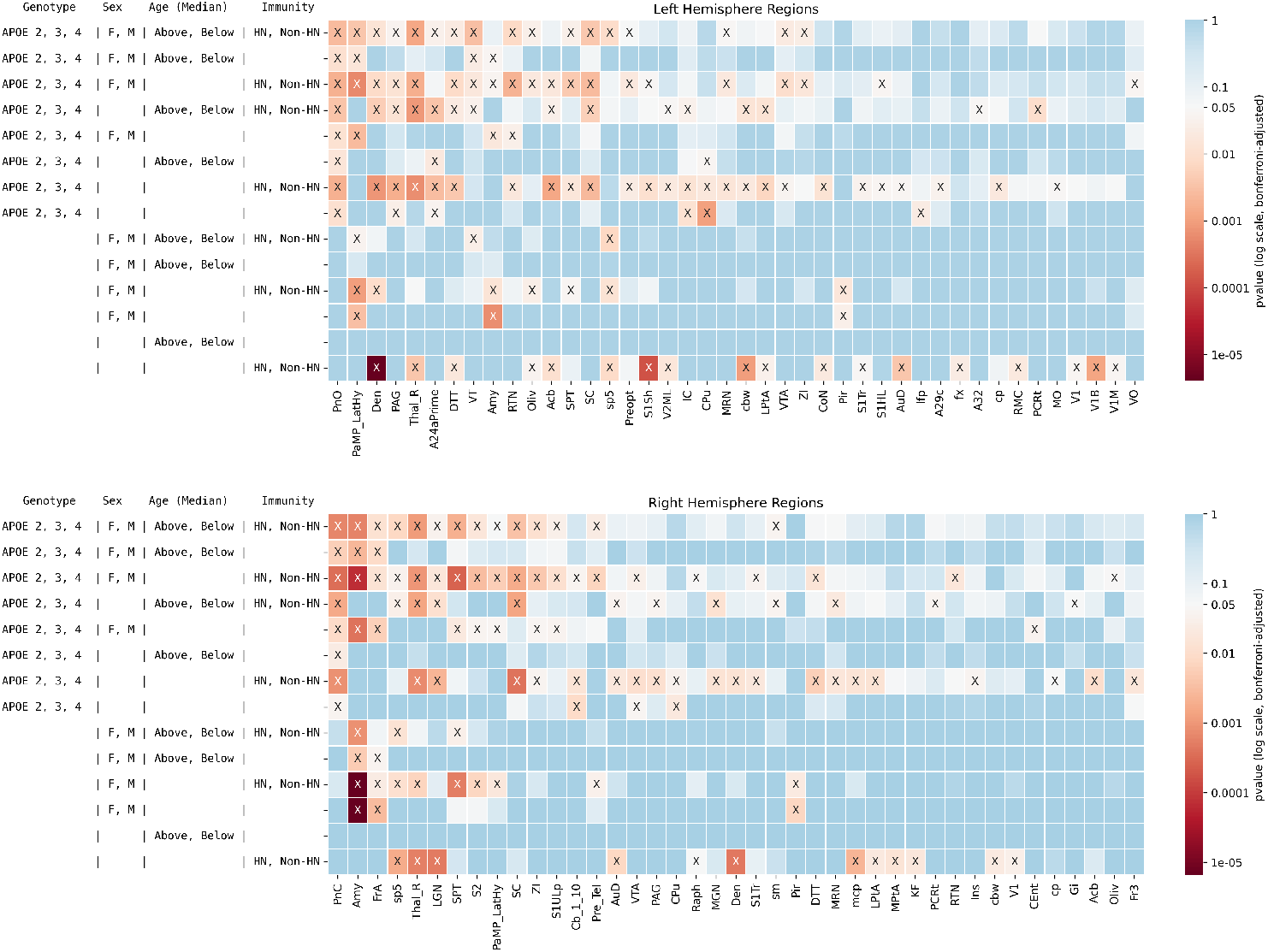
Significant Brain Regions From Testing for Effects of Genotypes, Sex, Age, and Innate Immunity. The results of tests conducted on each hemisphere are shown on top panel for left hemisphere and bottom panel for the right hemisphere. Significant results are shown by X. Brain regions in which no tests were significant are not shown for visualization purposes. A total of *n* = 114 mice were included in the tests. In all, 207 combinations of genotypes and factors were tested.

1. Age: the median age was not a pivotal factor, as age alone did not show any significant results for our cohort.
2. Sex: Significant sex-related differences were notable in regions traditionally associated with hormonal influence, such as the bilateral amygdala and left lateral hypothalamus; also the bilateral piriform cortex; right frontal association cortex. Three regions showed differences in each hemisphere.
3. APOE Genotype: The genotype factor showed a diverse influence on brain regions including the bilateral caudate putamen (CPu); left pons (PN, oral part), periaqueductal gray (PAG), cingulate cortex area 24a (A24a),inferior colliculus (IC), and longitudinal fasciculus of pons (lfp); right ventral tegmental area (VTA), and cerebellar nuclei. 6 regions in the left hemipshere and 9 in the right were affected.
4. Immunity: The contrast between mouse and humanized immune system mice yielded the largest number of significant results; and involved the bilateral dentate nucleus of the cerebellum, thalamus, parietal association cortex, auditory and visual cortex (V1B, V1 motor); left cochlear nucleus, fornix, cerebellar white matter, dorsal tenia tecta; and right raphe nuclei. 18 regions in the left hemisphere and 12 in the right were affected.

We performed the same analyses within each of the APOE genotypes Figure 4.

**Figure 4:**
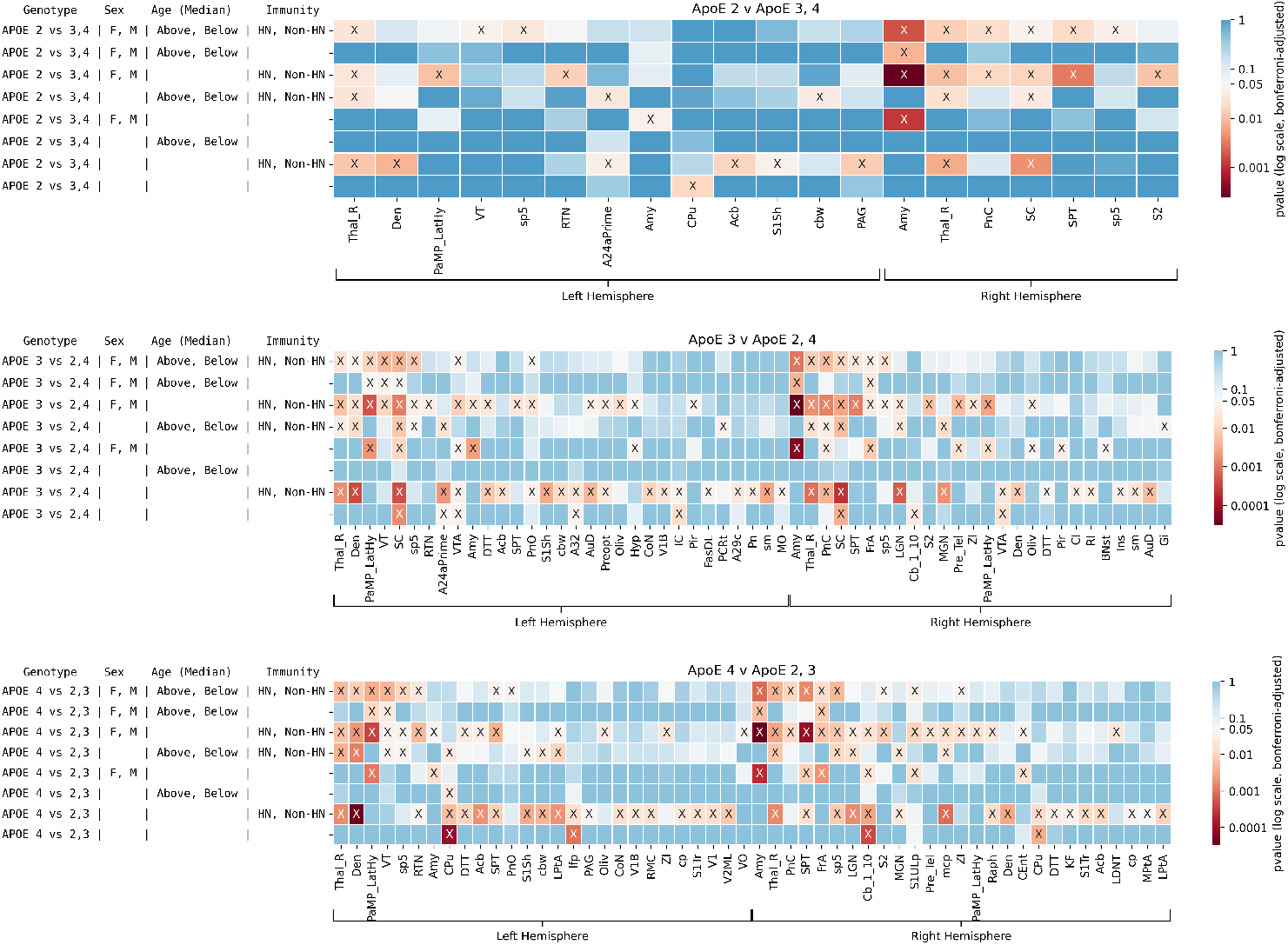
Significant Brain Regions From Testing for Effects of Sex, Age, and Immunity, for Each Major APOE Alelle. The results of tests conducted on different genotypes are shown in each panel. Significant results are shown by X. Brain regions in which no tests were significant are not shown for visualization purposes. A total of *n* = 114 mice were included in the tests.

In conclusion for this analysis in animals on a control diet, among all comparisons, the largest number of regions found to differ significantly was for immunity (18 on the left, 12 on the right hemisphere), followed by APOE allele (6 left, 4 on the right hemisphere), and then sex (3 regions in each hemisphere). A prominent role was found for the right amygdala and thalamus rest, as these regions were highly significant for 8 of the 15 comparisons for all factors. Interestingly, the pons were also significant in 8 of the comparisons but p values were higher. We noted that the interaction of APOE and immunity revealed a large number of regions (26 left hemisphere, 21 in the right hemisphere); as well as the interaction of APOE, sex and immunity (20 in each hemisphere). The combination of all factors revealed 17 regions in the left, and 14 in the right hemisphere, including the bilateral amygdala, thalamus and superior colliculus; right septum, and left dentatecerebellar nucleus.

When comparing ApoE2 versus ApoE3-4 mice, much fewer regions were significantly different, however we noted a role for the caudate putamen for genotypes comparisons; and for the amygdala when examining the role of sex, and the interaction of sex with immunity, where additional regions plaedy a role including septum, S2, hypothalamus, and thalamus including its reticular nucleus. Amygdala was present in the interaction of all factors, as well as septum, superior colliculus, and thalamus within APOE2 mice.

When comparing APOE3 mice versus the other strains we found a role for the superior colliculus, cingulate cortex, ventral tegmental area, and cerebellar nuclei for genotype comparisons. The largest differences in terms of the number of regions appeared for the effect of immunity (34 regions). The interaction of all factors included the thalamus, superior colliculus, sp5, pons bilaterally, as well as amygdala, septum, frontal association cortex, VTA, lateral hypothalamus, and dentate nucleus of cerebellum. Sex differences included amygdala, hypothalamus, bed nucleus of stria terminalis, but also frontal association cortex.

The caudate putamen, the cerebellum and longitudinal fasciculus of pons appeared as significant when comparing within the APOE4 vs other strains. Age was important within the APOE4 genotype, and pointed to a role for the caudate putamen. 36 regions were affected by immunity. For the interaction of all factors we noted the thalamus rest, septum, pons, and spinal trigemnial nerve bilaterally; as well as frontal association cortex, S1, S2, thalamic nuclei (reticular, ventral and zona incerta), lateral hypothalamus, amygdala, S1 and S2. Interestingly sex differences were associated with the entorhinal cortex.

In summary, certain regions that are typically implicated in Alzheimer’s disease, such as amygdala and thalamus were significantly different in all the tests. The effect of immunity in terms of the number of significant regions increased from APOE2 to APOE3 to APOE4.

### 3.3 Effect of Diet

Studying the effects of a high-fat diet (HFD) in ApoE mice is crucial for understanding the interplay between diet, APOE genotype, and neurodegenerative diseases, such as Alzheimer’s disease Figure 5.

**Figure 5:**
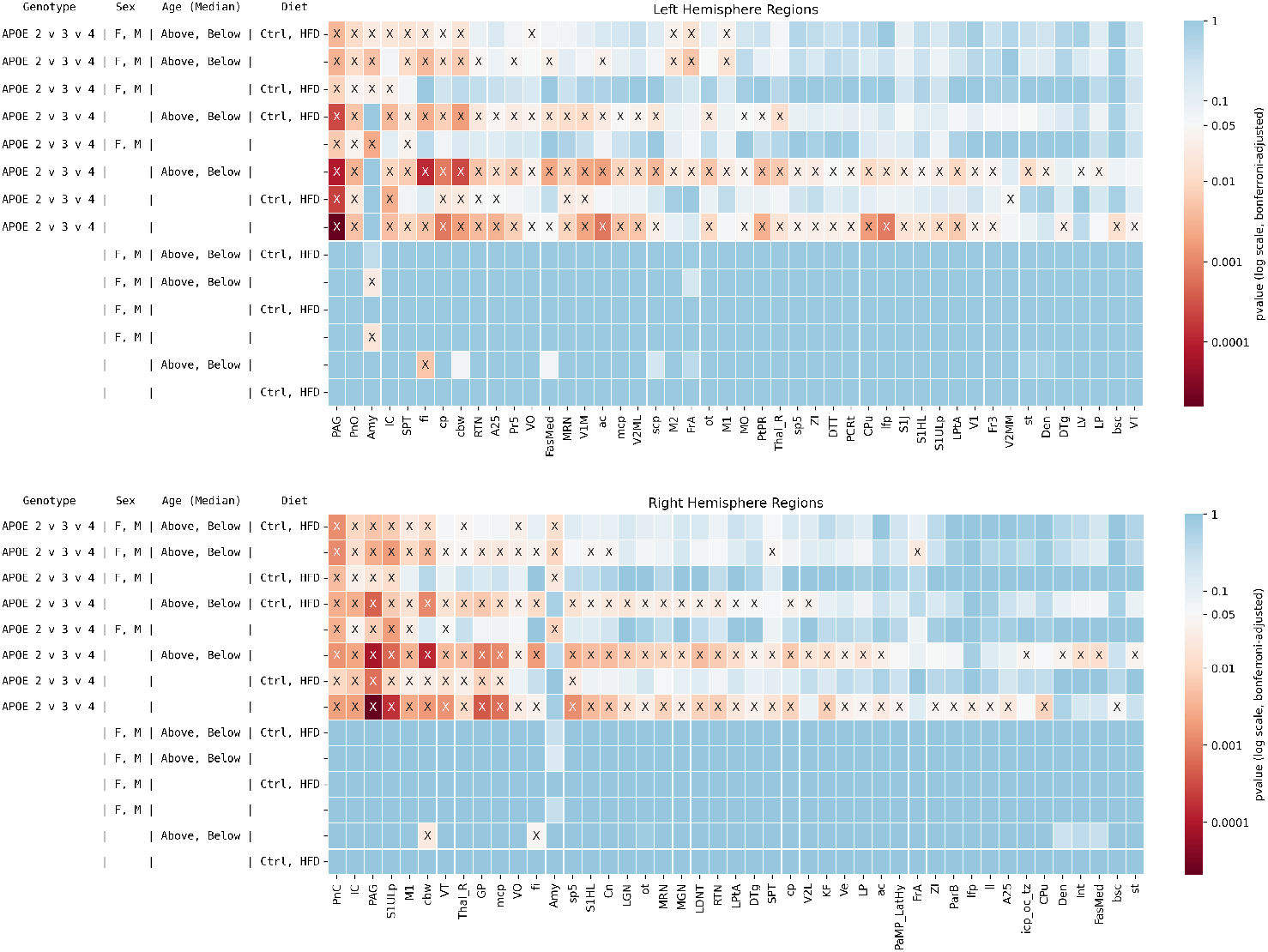
Significant Brain Regions From Testing for Effects of Genotypes, Sex, Age, and Diet. The results of tests conducted on each hemisphere are shown on top panel for left hemisphere and bottom panel for the right hemisphere. Significant results are shown by X. Brain regions in which no tests were significant are not shown for visualization purposes. A total of *n* = 75 mice were included in the tests. In all, 154 combinations of genotypes and factors were tested.

1. Diet: The diet treatment did not have an effect on its own, but did show significant interactions with ApoE genotypes, but not with age and sex. APOE by diet interaction showed an effect on the left pons, inferior colliculus, S1, cingulate cortex (A25), V1 amd V2, and midbrain reticular nuclei, cerebral peduncle and cerebellar white matter, right M1, thalamus, and globus pallidus.
2. Sex: Similarly, sex only showed a direct effect on one region, the amygdala (left), and showed significant interactions with ApoE genotypes for the bilateral amygdala and periaqueductal gray, left septum; right S1, M1, and inferior colliculus,
3. Age: Age had a significant effect for fimbria (bilateral), and right cerebellar white matter; and showed significant interactions with ApoE genotypes for 41 regions in the left hemisphere and 35 regions in the right hemisphere including V1, V2, S1, septum, caudate putamen, cerebellar nuclei, fimbria, and other major white matter tracts, as well as for the lateral ventricles.
4. Genotype: The APOE genotype factor showed a widespread influence on brain regions, including the bilateral periaqueductal gray (PAG), pons, caudate putamen; right globus palidus, left parietal association cortex, right lateral hypithalamus. Diet interacted with genotype for the bilateral PAG, pons, inferior coliculus, V2, thalamus (ventral), and also left cingulate cortex, and V1; right S1, M1, globus pallidus. The amygdala, M1, inferior colliculus, and PAG were present at the intersection of all factors; as well as left septum and fimbria; and right ventral orbital cortex.

Within group analyses showed differences in APOE2 and APOE4 mice. APOE3 differed less relative to the other two genotypes and only for APOE by sex by age in the amygdala and frontal association cortex. APOE by diet showed a role for the PAG and the logitudinal fasculucls of pons for the APOE4 comparison, while the APOE2 comparison showed differences in 34 regions, including septum.

For APOE4 the interaction of APOE by Age by Diet showed a role for PAG, IC, fimbria, optic tract, cerebral peduncle, cerebellar white matter bilaterally, then unilaterally for midbrain reticular nuclei, M1, V2, pons and white matter tracts including fimbria, cerebral peduncle, optic tracts, spinal trigeminal tract, cerebellar white matter bilaterally.

## 4 Discussion

Given the complexity of risk factors for LOAD, we have examined jointly changes in volume and FA covariances in aging mice modeling genetic risk for LOAD, as well as the effects of sex, diet and immunity. Our qualitative examination of the average connectomes support an important role of microstructural features such as FA, which may be an important feature to add to the battery of imaging parameters used to predict and follow the course of AD, and in addition to following the more widely studied regional brain atrophy patterns along the lifespan[44].

**Figure 6:**
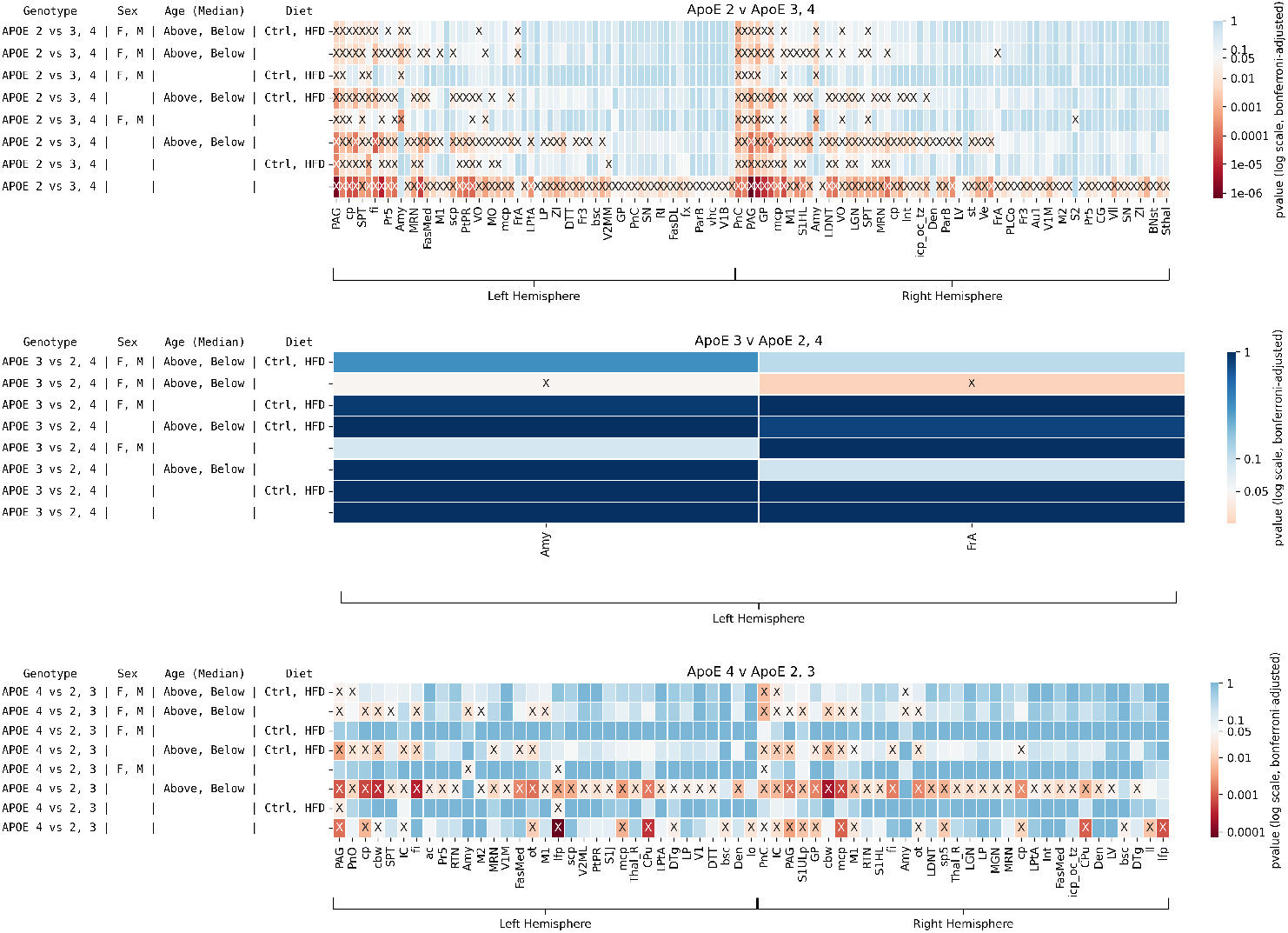
Significant Brain Regions From Testing for Effects of Sex, Age, and Diet Separated by Pairs of Genotypes. The results of tests conducted on different pair of genotypes are shown in each panel. Significant results are shown by X. Brain regions in which no tests were significant are not shown for visualization purposes. A total of *n* = 75 mice were included in the tests. Clearly, there are less differences between ApoE2 and ApoE3-4 group as much fewer regions were significantly different. Interestingly, certain regions that are typically implicated in Alzheimer’s disease, such as amygdala and thalamus show as significantly different in all the tests.

When studying animals on a regular diet we found that APOE genotype influenced primarily the network integration of the caudate putamen [45], but also the cingulate cortex [46] involved in mediating cognitive influences on emotion, response to threats, ventral tegmental area involved in regulating reward, learning, memory and addiction behaviors [47]; pons whose volume loss has been associated with greater neocortical amyloid burden [48], periaqueductal gray invoved in modulation of pain perception, and its memory, subsequently linked to anxiety and depression [49], inferior colliculus modulating auditory signal integration, and helping map physical space with both auditory and visual information [50]. These results support that APOE is involved in modulating brain networks architecture in regions known to be involved in learning, memory and emotion, functions impacted in AD, regions involved in reward processing and psychiatric conditions, such as depression, and with sensory function.

Sex differences consistently pointed primarily to a role of the amygdala, lateral hypothalamus, well known sexually dimorphic structures [51], as well as the piriform cortex, and frontal association cortex. Immunity effects were seen for a large number of regions, including the thalamus, lateral parietal association cortex, auditory and visual cortex, cerebellar white matter and dentate nuclei, but also fornix, the primary patway of the limbic system.

In terms of common findings for all APOE genotypes, the amygdala played an important role for sex differences, sex x immunity, sex x age, and for sex x age x immunity; and the thalamus for immunity, age and immunity, sex and immunity and the intersection of all factors.

The specific effect of APOE2 was important for the caudate putamen (CPu), and for this group, immunity identified 8 regions, pointing to a role for the thalamus bilaterally, as well as superior colliculus, accumbens, S1, cingulate cortex, and dentate cerebellar nucleus, predominantly for the left hemisphere. Sex differences isolated the amygdala. In addition to the amygdala, the interaction between sex and immunity pointed to a role for the septum. Age and immunity pointed to a role for the thalamus, bilaterally, and also cingulate cortex and superior colliculus. The intersection of all risk factors identified changes in 9 regions, underlining the bilateral amygdala, thalamus, and spinal trigeminal tract; and right septum, superior colliculus, pons.

The presence of APOE3 indicated additional regions (15 total) changed withe the combination of risk factors, besides the regions found for APOE2. These included the frontal association cortex, the lateral hypothalamus, and the dentate cerebellar nuclei. Immunity identified 34 changed regions, including the claustrum and insula.

The presence of APOE4 identified 17 regions where affected by all risk factors, adding the septum, S1 and S2; while 36 were affected by immunity, including the accumbens, septum S1, V1, V2 and parietal association cortices. Age was associated with changes to the caudate putamen for APOE4 only. APOE4 appeared to be most sensitive to interactions for age, sex, and immunity for animals on a regular diet, and in particular for immunity.

When studying animals exposed to control and high fat diet, the different APOE alleles affected 37 regions, up from the 4 and 6 regions in the previous analysis with the most significant changes for the periaqueductal gray, which has been previously involved in consummatory behaviors, and responses to reward, being proposed to mediate behaviors related to food consumption [52], where the bed nucleus of stria terminalis (BNST) and lateral hypothalamic GABAergic projections to the periaqueductal gray (PAG) may regulate feeding [53]. The interaction between APOE and HFD affected around 10 regions, the PAG, pons [54], and inferior colliculus which has been shown to respond to high levels of circulating glucose [55], and interestingly the cerebellar white matter, which has been show to have a high inflammatory response to a high fat diet [56], and was also prominent in the interaction of APOE x age, diet. This interaction also affected the fimbria, suggesting alterations in the memory processing system. The interaction of APOE, sex, and diet revealed a small set of regions including periaquductal gray (PAG), pons, amygdala and inferior colliculus, primary somatosesory cortex (S1), and fimbria. The interaction of all risk factors included the amygdala, similarly to our previous analyses, but also the PAG, motor cortex, and fimbria.

Interestingly genotype specific changes were more pronounced in APOE2 and APOE4 mice, than in APOE3 mice, indicating resilience of APOE3 mice to nework changes following a high fat diet. The APOE2 effect was present in a large number of regions (>70), including the bed nucleus of stria terminalis (BNST) periaquductal gray (PAG), and plobus pallidus (GP), associated with alcohol and opiate abuse [57]. Interestingly the interaction with sex was also affecting 11 regions, including septum and amygdala. The effect of APOE4 was present in 25 regions, and the interaction with aging caused changes in 58 regions.

### 4.1 Comparison with Status of the Field

While most studies so far examine one biomarker at a time, either atrophy of texture such as fractional anisotropy changes in AD or animal models of AD, here we propose a join approach for examining the signal in volume and texture subject specific covariance networks. Our methods rely on joint embedding of connectomes, and distance correlation metrics. Other methods for joint signal analyses have used multi canonical correlation [58][59][60], similarity driven multiview embeddings [61], SVM [62], and complex deep learning models [63] that an integrate multi MRI as well as clinical [64] [65] and omics data. The advent of novel methods for mulitvariate integrative modeling is a promising strategy for mining multi dimensional data sets, which then serve to better understand complex, multifactorial neurodegenerative diseases. Our methods strengthened the body of literature describing mouse models of genetic risk for AD to examine single subject networks that revealed vulnerable nodes to multiple risk factors, underlining the network integration changes for the caudate putamen, amygdala and thalamus in these mouse models. Our results support widespread white matter changes due to exposure to a high fat diet, further supporting network disease propagation models.

### 4.2 Limitations

Interestingly, age was not a pivotal factor in our analysis, supporting that its effects were uniform and widespread throughout the brain, but we acknowledge that we had a narrow age span. Future studies can examine younger mice and a wider age span may better reveal the interactions of age with APOE and other risk factors. Our study is limited in that we could only study the effect of diet in the APOExHN lines, and had reduced power. Also, mice do not perfectly replicate the complexities of human AD genetic risk interactions and the heterogeneity of human populations. Additionally, studying sex differences would benefit from using ovariectomized female mice to reveal menopause associated effects.

### 4.3 Conclusion

We present the application of novel models that integrate multivariate data to account for changes in multiple imaging based parameters. Our study in mouse models of genetic risk for LOAD revealed a role for the amygdala, caudate putamen in normally aging mice, widespread effects of innate immunity and more expansive, genotype specific changes in response to a high fat diet than under control conditions

## 5 Code

The code and data for these analyses are available at GitHub repository at https://github.com/neurodata/alzheimers-mouse. The analysis relied on graspologic [66], NumPy [67], SciPy [68], Pandas [69], and NetworkX [70]. Plotting was performed using matplotlib [71] and Seaborn [72].

### Author Contributions

EWB, JC, and AB wrote the manuscript; EWB, JC, JTV, and AB designed the statistical methods and study conception/design; EWB and JC performed the statistical analyses and data post-processing; RJA, AM, JAS, HSM, ZYH, and AB collected the data and performed all data pre-processing; all authors contributed valuable feedback and discussion throughout the work.

## Acknowledgements

The authors are grateful for the support from the National Institute of Health (NIH) through the National Institute on Aging (NIA) via Research Projects RF1AG057895, R01AG066184, and R01AG070149.

## Notes

https://github.com/neurodata/alzheimers-mouse

### Competing Interest Statement

The authors have declared no competing interest.

## References

[1] EH Corder, AM Saunders, WJ Strittmatter, D. Schmechel, PC Gaskell, GW Small, AD Roses, JL Haines, and MA Pericak-Vance. Gene dose of apolipoprotein e type 4 allele and the risk of alzheimer’s disease in late-onset families. Science, 1993.

[2] Lilian Calderón-Garcidueñas, Ricardo Torres-Jardón, Randy J Kulesza, Yusra Mansour, Luis Oscar González-González, Angélica Gónzalez-Maciel, Rafael Reynoso-Robles, and Partha S Mukherjee. Alzheimer disease starts in childhood in polluted metropolitan mexico city. a major health crisis in progress. Environmental research, 183:109137, 2020.

[3] Sam Norton, Fiona E Matthews, Deborah E Barnes, Kristine Yaffe, and Carol Brayne. Potential for primary prevention of alzheimer’s disease: an analysis of population-based data. Lancet Neurology, 2014.

[4] Ron Brookmeyer, S Gray, and C Kawas. Projections of alzheimer’s disease in the united states and the public health impact of delaying disease onset. American Journal of Public Health, 1998.

[5] Clive Ballard, Serge Gauthier, Anne Corbett, Carol Brayne, Dag Aarsland, and Emma Jones. Alzheimer’s disease. Lancet, 2011.

[6] Ashford J Wesson. Apoe genotype effects on alzheimer’s disease onset and epidemiology. J Mol Neurosci, 23(3):157–65, 2004.

[7] David R. Riddell, Hua Zhou, Kevin Atchison, Helen K. Warwick, Peter J. Atkinson, Julius Jefferson, Lin Xu, Suzan Aschmies, Yolanda Kirksey, Yun Hu, Erik Wagner, Adrienne Parratt, Jane Xu, Zhuting Li, Margaret M. Zaleska, J. Steve Jacobsen, Menelas N. Pangalos, and Peter H. Reinhart. Impact of apolipoprotein e (apoe) polymorphism on brain apoe levels. Journal of Neuroscience, 28(45):11445 11453, 2008. doi: 10.1523/JNEUROSCI.1972-08.2008. URL https://www.scopus.com/inward/record.uri?eid=2-s2.0-58149251844&doi=10.1523%2fJNEUROSCI.1972-08.2008&partnerID=40&md5=3752cb5062536406a84615d232917e49. Cited by: 266; All Open Access, Green Open Access, Hybrid Gold Open Access.

[8] Yaqiong Xiao, Jiaojian Wang, Kaiyu Huang, Lei Gao, Shun Yao, Alzheimers Disease Neuroimaging Initiative, et al. Progressive structural and covariance connectivity abnormalities in patients with alzheimers disease. Frontiers in Aging Neuroscience, 14:1064667, 2023.

[9] Anne Hafkemeijer, Christiane Möller, Elise GP Dopper, Lize C Jiskoot, Annette A van den Berg-Huysmans, John C van Swieten, Wiesje M van der Flier, Hugo Vrenken, Yolande AL Pijnenburg, Frederik Barkhof, et al. Differences in structural covariance brain networks between behavioral variant frontotemporal dementia and alzheimer’s disease. Human brain mapping, 37(3):978–988, 2016.

[10] Sambit Panda, Cencheng Shen, Ronan Perry, Jelle Zorn, Antoine Lutz, Carey E Priebe, and Joshua T Vogelstein. High-dimensional and universally consistent k-sample tests. arXiv preprint 1910.08883, 2019.

[11] Alexandra Badea and G Allan Johnson. Magnetic resonance microscopy. Stud Health Technol Inform, 185:153–84, 2013.

[12] Alexandra Badea, Alaa Kamnaksh, Robert J Anderson, Evan Calabrese, Joseph B Long, and Denes V Agoston. Repeated mild blast exposure in young adult rats results in dynamic and persistent microstructural changes in the brain. NeuroImage: Clinical, 18:60–73, 2018.

[13] Robert W Mahley, Karl H Weisgraber, and Yadong Huang. Apolipoprotein e4: a causative factor and therapeutic target in neuropathology, including alzheimers disease. Proceedings of the National Academy of Sciences, 103(15):5644–5651, 2006.

[14] Yu-Wen Alvin Huang, Bo Zhou, Amber M Nabet, Marius Wernig, and Thomas C Südhof. Differential signaling mediated by apoe2, apoe3, and apoe4 in human neurons parallels alzheimer’s disease risk. Journal of Neuroscience, 39(37):7408–7427, 2019.

[15] Thomas J Gross, Karol Kremens, Linda S Powers, Brandi Brink, Tina Knutson, Frederick E Domann, Robert A Philibert, Mohammed M Milhem, and Martha M Monick. Epigenetic silencing of the human nos2 gene: rethinking the role of nitric oxide in human macrophage inflammatory responses. The Journal of Immunology, 192(5):2326–2338, 2014.

[16] Michael D Hoos, Michael P Vitek, Lisa A Ridnour, Joan Wilson, Marilyn Jansen, Angela Everhart, David A Wink, and Carol A Colton. The impact of human and mouse differences in nos2 gene expression on the brains redox and immune environment. Molecular neurodegeneration, 9:1–15, 2014.

[17] Hae Sol Moon, Ali Mahzarnia, Jacques Stout, Robert J. Anderson, Madison Strain, Jessica T. Tremblay, Zay Yar Han, Andrei Niculescu, Anna MacFarlane, Jasmine King, Allison Ashley-Koch, Darin Clark, Michael W. Lutz, and Alexandra Badea. Multivariate investigation of aging in mouse models expressing the alzheimers protective apoe2 allele: integrating cognitive metrics, brain imaging, and blood transcriptomics, December 2023. URL http://dx.doi.org/10.1007/s00429-023-02731-x.

[18] Alexandra Badea, Wenlin Wu, Jordan Shuff, Michele Wang, Robert J Anderson, Yi Qi, G Allan Johnson, Joan G Wilson, Serge Koudoro, Eleftherios Garyfallidis, et al. Identifying vulnerable brain networks in mouse models of genetic risk factors for late onset alzheimers disease. Frontiers in neuroinformatics, 13:72, 2019.

[19] Jonathan I Tamir, Frank Ong, Joseph Y Cheng, Martin Uecker, and Michael Lustig. Generalized magnetic resonance image reconstruction using the berkeley advanced reconstruction toolbox. In ISMRM Workshop on Data Sampling & Image Reconstruction, Sedona, AZ, 2016.

[20] Nian Wang, Robert J Anderson, Alexandra Badea, Gary Cofer, Russell Dibb, Yi Qi, and G Allan Johnson. Whole mouse brain structural connectomics using magnetic resonance histology. Brain Structure and Function, 223:4323–4335, 2018.

[21] J-Donald Tournier, Fernando Calamante, and Alan Connelly. Mrtrix: diffusion tractography in crossing fiber regions. International journal of imaging systems and technology, 22(1):53–66, 2012.

[22] Robert James Anderson, Nian Wang, James J Cook, Gary P Cofer, Russell Dibb, G Allan Johnson, and Alexandra Badea. A high performance computing cluster implementation of compressed sensing reconstruction for mr histology. In Proc. Intl. Soc. Mag. Reson. Med, volume 26, 2018.

[23] Robert J Anderson, James J Cook, Natalie Delpratt, John C Nouls, Bin Gu, James O McNamara, Brian B Avants, G Allan Johnson, and Alexandra Badea. Small animal multivariate brain analysis (samba)–a high throughput pipeline with a validation framework. Neuroinformatics, 17:451–472, 2019.

[24] Evan Calabrese, Alexandra Badea, Gary Cofer, Yi Qi, and G Allan Johnson. A diffusion mri tractography connectome of the mouse brain and comparison with neuronal tracer data. Cerebral cortex, 25(11):4628–4637, 2015.

[25] Steven Winter, Ali Mahzarnia, Robert J Anderson, Zay Y Han, Jessica J Tremblay, Daniel Marcellino, David Dunson, and Alexandra Badea. Apoe, immune factors, sex, and diet interact to shape brain networks in mouse models of aging. bioRxiv, pages 2023–10, 2023.

[26] Alexandra Badea. Mouse brain atlas ex vivo mri, February 2024. URL 10.5281/zenodo.10652239.

[27] Stephen J. Young and Edward R. Scheinerman. Random Dot Product Graph Models for Social Networks. In Anthony Bonato and Fan R. K. Chung, editors, Algorithms and Models for the Web-Graph, Lecture Notes in Computer Science, pages 138–149. Springer Berlin Heidelberg, 2007. ISBN 978-3-540-77004-6.

[28] Daniel L. Sussman, Minh Tang, Donniell E. Fishkind, and Carey E. Priebe. A Consistent Adjacency Spectral Embedding for Stochastic Blockmodel Graphs. Journal of the American Statistical Association, 107(499):1119–1128, September 2012. ISSN 0162-1459. doi: 10.1080/01621459.2012.699795. URL https://amstat.tandfonline.com/doi/full/10.1080/01621459.2012.699795.

[29] Minh Tang, Avanti Athreya, Daniel L. Sussman, Vince Lyzinski, and Carey E. Priebe. A nonparametric two-sample hypothesis testing problem for random graphs. Bernoulli, 23(3):1599–1630, August 2017. ISSN 1350-7265. doi: 10.3150/15-BEJ789. URL https://projecteuclid.org/euclid.bj/1489737619.

[30] Mu Zhu and Ali Ghodsi. Automatic dimensionality selection from the scree plot via the use of profile likelihood. Computational Statistics & Data Analysis, 51(2):918–930, 2006.

[31] Ronald A Fisher. Xv.the correlation between relatives on the supposition of mendelian inheritance. Earth and Environmental Science Transactions of the Royal Society of Edinburgh, 52(2):399–433, 1919.

[32] Maurice S Bartlett. Multivariate analysis. Supplement to the journal of the royal statistical society, 9(2):176–197, 1947.

[33] Frank Yates. The analysis of multiple classifications with unequal numbers in the different classes. Journal of the American Statistical Association, 29(185):51–66, 1934.

[34] Keenan A Pituch and James P Stevens. Applied Multivariate Statistics for the Social Sciences. Taylor & Francis, December 2015. URL https://www.taylorfrancis.com/books/edit/10.4324/9781315814919/applied-multivariate-statistics-social-sciences-james-stevens-keenan-pituch.

[35] Russell Warne. A primer on multivariate analysis of variance (manova) for behavioral scientists. Practical Assessment, Research, and Evaluation, 19(1):17, 2014.

[36] Gábor J. Székely, Maria L. Rizzo, and Nail K. Bakirov. Measuring and testing dependence by correlation of distances. The Annals of Statistics, 35(6):2769–2794, December 2007. ISSN 0090-5364. doi: 10.1214/009053607000000505.

[37] Avanti Athreya, Donniell E. Fishkind, Minh Tang, Carey E. Priebe, Youngser Park, Joshua T. Vogelstein, Keith Levin, Vince Lyzinski, Yichen Qin, and Daniel L. Sussman. Statistical Inference on Random Dot Product Graphs: a Survey. Journal of Machine Learning Research, 18(226):1–92, 2018. URL http://jmlr.org/papers/v18/17-448.html.

[38] K. Levin, A. Athreya, M. Tang, V. Lyzinski, and C. E. Priebe. A Central Limit Theorem for an Omnibus Embedding of Multiple Random Dot Product Graphs. In 2017 IEEE International Conference on Data Mining Workshops (ICDMW), pages 964–967, November 2017. doi:10.1109/ICDMW.2017.132.

[39] Daniel L. Sussman, Minh Tang, Donniell E. Fishkind, and Carey E. Priebe. A consistent adjacency spectral embedding for stochastic blockmodel graphs. Journal of the American Statistical Association, 107(499):1119–1128, 2012. doi: 10.1080/01621459.2012.699795. URL https://doi.org/10.1080/01621459.2012.699795.

[40] Daniel L. Sussman, Minh Tang, and Carey E. Priebe. Consistent latent position estimation and vertex classification for random dot product graphs. IEEE Transactions on Pattern Analysis and Machine Intelligence, 36:48–57, 2014. doi: 10.1109/TPAMI.2013.135.

[41] Vince Lyzinski, Daniel L. Sussman, Minh Tang, Avanti Athreya, and Carey E. Priebe. Perfect clustering for stochastic blockmodel graphs via adjacency spectral embedding. Electronic Journal of Statistics, 8(2):2905–2922, 2014. doi: 10.1214/14-EJS978. URL https://doi.org/10.1214/14-EJS978.

[42] Cencheng Shen, Sambit Panda, and Joshua T. Vogelstein. The Chi-Square Test of Distance Correlation. Journal of Computational and Graphical Statistics, January 2022. ISSN 1061-8600. doi: 10.1080/10618600.2021.1938585.

[43] Sture Holm. A simple sequentially rejective multiple test procedure. Scandinavian journal of statistics, pages 65–70, 1979.

[44] Milton Camacho, Matthias Wilms, Hannes Almgren, Kimberly Amador, Richard Camicioli, Zahinoor Ismail, Oury Monchi, and Nils D. Forkert. Exploiting macro- and micro-structural brain changes for improved Parkinson’s disease classification from MRI data. npj Parkinson’s Disease, 10(43):1–12, February 2024. ISSN 2373-8057. doi: 10.1038/s41531-024-00647-9.

[45] Margaret Caroline Stapleton, Stefan Paul Koch, Devin Raine Everaldo Cortes, Samuel Wyman, Kristina E Schwab, Susanne Mueller, Christopher Gordon McKennan, Philipp Boehm-Sturm, and Yijen Lin Wu. Apolipoprotein-e deficiency leads to brain network alteration characterized by diffusion mri and graph theory. Frontiers in Neuroscience, 17:1183312, 2023.

[46] Eric M Reiman, Kewei Chen, Gene E Alexander, Richard J Caselli, Daniel Bandy, David Osborne, Ann M Saunders, and John Hardy. Correlations between apolipoprotein e ε4 gene dose and brain-imaging measurements of regional hypometabolism. Proceedings of the National Academy of Sciences, 102(23):8299–8302, 2005.

[47] J Cai and Q Tong. Anatomy and function of ventral tegmental area glutamate neurons. front neural circuits 16: 867053, 2022.

[48] Heidi IL Jacobs, Adrienne ODonnell, Claudia L Satizabal, Cristina Lois, Daniel Kojis, Bernard J Hanseeuw, Emma Thibault, Justin S Sanchez, Rachel F Buckley, Qiong Yang, et al. Associations between brainstem volume and alzheimers disease pathology in middle-aged individuals of the framingham heart study. Journal of Alzheimer’s Disease, 86(4):1603–1609, 2022.

[49] Miriam Mokhtar and Paramvir Singh. Neuroanatomy, periaqueductal gray. 2020.

[50] Iori Ito, Rose Chik-Ying Ong, Baranidharan Raman, and Mark Stopfer. Sparse odor representation and olfactory learning. Nat. Neurosci., 11(10):1177–1184, October 2008.

[51] Jill M Goldstein, Larry J Seidman, Nicholas J Horton, Nikos Makris, David N Kennedy, Verne S Caviness Jr, Stephen V Faraone, and Ming T Tsuang. Normal sexual dimorphism of the adult human brain assessed by in vivo magnetic resonance imaging. Cerebral cortex, 11(6):490–497, 2001.

[52] Valerie Lee Tryon and Sheri JY Mizumori. A novel role for the periaqueductal gray in consummatory behavior. Frontiers in behavioral neuroscience, 12:178, 2018.

[53] Sijia Hao, Hongbin Yang, Xiaomeng Wang, Yang He, Haifeng Xu, Xiaotong Wu, Libiao Pan, Yijun Liu, Huifang Lou, Han Xu, et al. The lateral hypothalamic and bnst gabaergic projections to the anterior ventrolateral periaqueductal gray regulate feeding. Cell reports, 28(3):616–624, 2019.

[54] Norio Harada and Nobuya Inagaki. Regulation of food intake by intestinal hormones in brain. Journal of Diabetes Investigation, 13(1):17, 2022.

[55] Olga N. Vasilyeva, Susan T. Frisina, Xiaoxia Zhu, Joseph P. Walton, and Robert D. Frisina. Interactions of hearing loss and diabetes mellitus in the middle age cba/caj mouse model of presbycusis. Hearing Research, 249(1):44–53, 2009. ISSN 0378-5955. doi: 10.1016/j.heares.2009.01.007. URL https://www.sciencedirect.com/science/article/pii/S0378595509000100.

[56] Owein Guillemot-Legris, Julien Masquelier, Amandine Everard, Patrice D Cani, Mireille Alhouayek, and Giulio G Muccioli. High-fat diet feeding differentially affects the development of inflammation in the central nervous system. Journal of neuroinflammation, 13:1–11, 2016.

[57] Nismat Javed and Marco Cascella. Neuroanatomy, globus pallidus. 2020.

[58] Brian B Avants, David J Libon, Katya Rascovsky, Ashley Boller, Corey T McMillan, Lauren Massimo, H Branch Coslett, Anjan Chatterjee, Rachel G Gross, and Murray Grossman. Sparse canonical correlation analysis relates network-level atrophy to multivariate cognitive measures in a neurodegenerative population. Neuroimage, 84:698–711, 2014.

[59] Alexandra Badea, Natalie A Delpratt, RJ Anderson, Russell Dibb, Yi Qi, Hongjiang Wei, Chunlei Liu, William C Wetsel, Brian B Avants, and Carol Colton. Multivariate mr biomarkers better predict cognitive dysfunction in mouse models of alzheimer’s disease. Magnetic resonance imaging, 60: 52–67, 2019.

[60] Ali Mahzarnia, Jacques A Stout, Robert J Anderson, Hae Sol Moon, Zay Yar Han, Kate Beck, Jeffrey N Browndyke, David B Dunson, Kim G Johnson, Richard J OBrien, et al. Identifying vulnerable brain networks associated with alzheimers disease risk. Cerebral Cortex, 33(9):5307–5322, 2023.

[61] Brian B. Avants, Nicholas J. Tustison, and James R. Stone. Similarity-driven multi-view embeddings from high-dimensional biomedical data. Nature computational science, 1(2):143, February 2021. doi: 10.1038/s43588-021-00029-8.

[62] Daoqiang Zhang and Dinggang Shen. Multi-modal multi-task learning for joint prediction of multiple regression and classification variables in alzheimer’s disease. NeuroImage, 59(2):895–907, 2012. ISSN 1053-8119. doi: 10.1016/j.neuroimage.2011.09.069. URL https://www.sciencedirect.com/science/article/pii/S105381191101144X.

[63] Alexander Khvostikov, Karim Aderghal, Andrey Krylov, Gwenaelle Catheline, and Jenny Benois-Pineau. 3d inception-based cnn with smri and md-dti data fusion for alzheimer’s disease diagnostics. arXiv preprint 1809.03972, 2018.

[64] Hae Sol Moon, Ali Mahzarnia, Jacques Stout, Robert J Anderson, Zay Yar Han, Cristian T Badea, and Alexandra Badea. Predicting brain age and associated structural networks in mouse models with humanized apoe alleles using integrative and interpretable graph neural networks. In Medical Imaging 2024: Computer-Aided Diagnosis, volume 12927, pages 268–275. SPIE, 2024.

[65] Hae Sol Moon, Ali Mahzarnia, Jacques Stout, Robert J Anderson, Cristian T Badea, and Alexandra Badea. Feature attention graph neural network for estimating brain age and identifying important neural connections in mouse models of genetic risk for alzheimer’s disease. bioRxiv, pages 2023–12, 2023.

[66] Jaewon Chung, Benjamin D Pedigo, Eric W Bridgeford, Bijan K Varjavand, Hayden S Helm, and Joshua T Vogelstein. GraSPy: Graph Statistics in Python. J. Mach. Learn. Res., 20(158):7, 2019.

[67] Charles R. Harris, K. Jarrod Millman, Stéfan J. van der Walt, Ralf Gommers, Pauli Virtanen, David Cournapeau, Eric Wieser, Julian Taylor, Sebastian Berg, Nathaniel J. Smith, Robert Kern, Matti Picus, Stephan Hoyer, Marten H. van Kerkwijk, Matthew Brett, Allan Haldane, Jaime Fernández del Río, Mark Wiebe, Pearu Peterson, Pierre Gérard-Marchant, Kevin Sheppard, Tyler Reddy, Warren Weckesser, Hameer Abbasi, Christoph Gohlke, and Travis E. Oliphant. Array programming with NumPy. Nature, 585(7825):357–362, September 2020. ISSN 1476-4687. doi: 10.1038/s41586-020-2649-2. URL https://www.nature.com/articles/s41586-020-2649-2. Number: 7825 Publisher: Nature Publishing Group.

[68] Pauli Virtanen, Ralf Gommers, Travis E. Oliphant, Matt Haberland, Tyler Reddy, David Cournapeau, Evgeni Burovski, Pearu Peterson, Warren Weckesser, Jonathan Bright, Stéfan J. van der Walt, Matthew Brett, Joshua Wilson, K. Jarrod Millman, Nikolay Mayorov, Andrew R. J. Nelson, Eric Jones, Robert Kern, Eric Larson, C. J. Carey, lhan Polat, Yu Feng, Eric W. Moore, Jake VanderPlas, Denis Laxalde, Josef Perktold, Robert Cimrman, Ian Henriksen, E. A. Quintero, Charles R. Harris, Anne M. Archibald, Antônio H. Ribeiro, Fabian Pedregosa, and Paul van Mulbregt. SciPy 1.0: fundamental algorithms for scientific computing in Python. Nature Methods, 17(3):261–272, March 2020. ISSN 1548-7105. doi: 10.1038/s41592-019-0686-2. URL https://www.nature.com/articles/s41592-019-0686-2. Number: 3 Publisher: Nature Publishing Group.

[69] Wes McKinney. pandas: a Foundational Python Library for Data Analysis and Statistics. Python for high performance and scientific computing, 14(9):1–9, 2011. Publisher: Seattle.

[70] Aric Hagberg, Pieter Swart, and Daniel Schult. Exploring network structure, dynamics, and function using networkx. Technical Report LA-UR-08-05495; LA-UR-08-5495, Los Alamos National Lab. (LANL), Los Alamos, NM (United States), January 2008. URL https://www.osti.gov/biblio/960616.

[71] John D. Hunter. Matplotlib: A 2D Graphics Environment. Computing in Science & Engineering, 9 (3):90–95, May 2007. ISSN 1558-366X. doi: 10.1109/MCSE.2007.55. Conference Name: Computing in Science & Engineering.

[72] Michael L. Waskom. seaborn: statistical data visualization. Journal of Open Source Software, 6 (60):3021, 2021. doi: 10.21105/joss.03021. URL https://doi.org/10.21105/joss.03021. Publisher: The Open Journal.

